# Distribution And Population Of Chukar Partrigde (*Alectoris chukar*) In District Bajuar, KPK, Pakistan

**DOI:** 10.1101/2021.11.05.467502

**Authors:** Durr e Shahwar, Atufa Kawan, Hina Mukhtar, Inam Ullah

**Affiliations:** Department of Forestry and Wildlife Management, University of Haripur KPK, Pakistan; College of Fisheries, Huazhong Agricultural University, Wuhan, China; College of Food Science and Technology, Huazhong Agricultural University, Wuhan, China

**Keywords:** *Alectoris chukar*, Chukar Partrigde, Bajaur, population, distribution

## Abstract

The Chukar Partridge (*Alectoris chukar*) is a wild bird of the Phasianidae family. In Pakistan, it occurs in a wide range of rugged, sloping and dry areas, rising to the interior of the Himalayas, the western Himalayas and plains, the higher piedmont valleys on the dry slopes of the Baluchistan provinces and Punjab and Sindh provinces. It is the “national bird” of Pakistan, however, little is known about its ecology and reproduction in the northern part of the country. This study investigated the existence, distribution and population of birds in the Bajaur region of Khyber Pakhtunkhwa (KP) Province from December 2019 to August 2020. Usually in the early morning (5 to 78 AM) and evening (4 to 9 pm). The areas occupied by *Alectoris chukar* mainly include *Mulberry, Ficus banyan, Gum arabic, Arabian acacia, Barberes lycium, Dilberia sisso, Melia azedarach, Ailanthus, Alalthus altissima, Zanthoxylum alatum, Olea europaea, Olea europaea, Ingres, Celtis eriocarpa* and *Pinus wallichiana*. Habitat destruction (overgrazing, mowing and landslides) caused by hunting, shooting, capture, explosives, excavation, and road construction is the main threat to existing wildlife (including *Alectoris chukar*).

## Introduction

A common game bird of the family Phasianidae is Chukar Partridge (*Alectoris chukar* Gray, 1830). Its origin in Asia includes Palestine, Lebanon, Turkey, Iran, Afghanistan, Pakistan and India, from the western Himalayas to Nepal (Birdlife International, 2016). It is usually found at mean sea level (amsl) between 2000 m and 4000 m above sea level, but it is also found at 600 m in Pakistan (Rasmussen and Anderton, 2005). Chukar can be found in the field in gatherings of 2-4 or typically 5-7 birds. From March, they begin to mix and raise the brood in the middle of early April to July, depending on the elevation. The breeding begins at low altitude right on time when compared to the higher rise (Roberts, 1991) that it travels to snow-topped fields in the Himalayas and does not begin to rise until late June. Roberts (1991) stated that Chukar nesting occurred over a long period of time, depending on altitude and latitude. Some birds in Himalayas ascend to alpine pastures and do not start breeding until late June. However, breeding can start early (from late March) at lower elevations. In the Salt Range, the main nesting season is April to May with a typical clutch size of 6 to 9 eggs, but the clutch size may range from 15 to 19 eggs in heavier rainfall areas.

Chukar Partridge is the national bird of Pakistan and a wild bird. However, in the country’s natural environment, there is insufficient scientific data on its ecology and breeding. The area Bajuar is contained within its local area, but there is no previous data record for this area. The current study aims to investigate the distribution and population of Chukar partridge in the Bajaur region of Khyber Pakhtunkhwa province.

## Methods

### Study Area

The current research has been carried out in the Bajaur area between 34.795100° latitude and 71.540594° longitude in the southeast. The Bajaur district was a semi-independent area before 1960 and was considered a place with inconvenient transportation. In 1960, Bajaur was declared a subdivision of the Malakand region, and in December 1973 it was declared a Federal Tribal Area (FATA). The Bajaur district is administratively divided into two districts, namely the Khar district and the Nawagai district and seven tehsils. The Bajaur district borders the Mohmand district in the southwest, and the Dir district borders Afghanistan in the northwest. The Bajaur region includes an arid subtropical area with harsh winters and hot summers. Winter begins in December and lasts until February, during which the temperature drops below zero. May to September is the warmest month of the year. The average annual rainfall is 800 mm /year. The terrain is mostly mountainous (Figure 1).

**Fig. 1:**
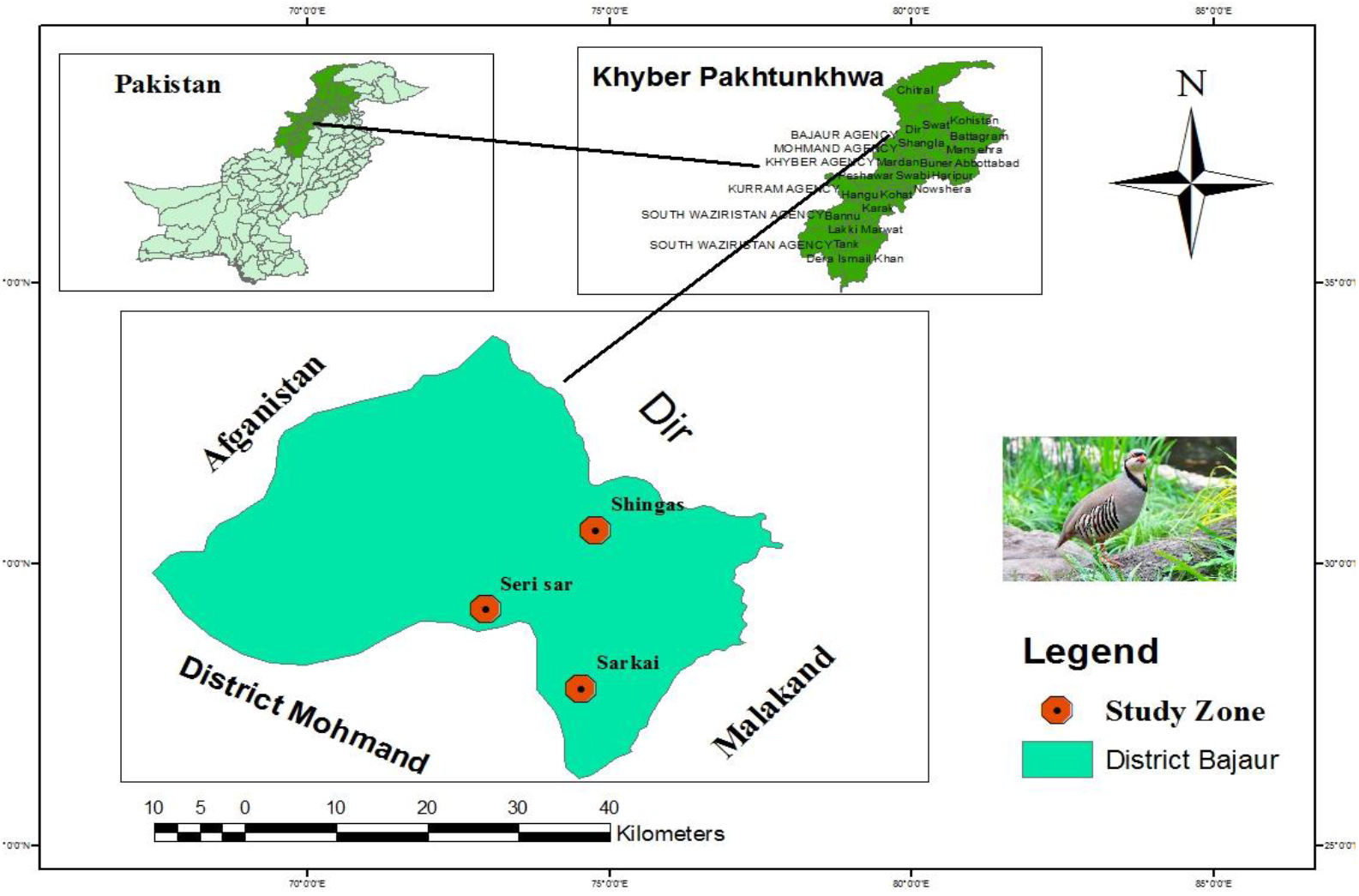
Map of the District Bajaur, Khyber Pakhtunkhwa, Pakistan

Nine months research survey was conducted from December 2019 to August 2020 to determine the distribution and population of *Alectoris chukar* in Bajaur district. To carry out the research area was divided into three main research zones, A, B and C (Figure 1; Table 1). During the investigation, direct and indirect methods were used to collect information about its distribution and population. Calls, feathers, feces and information collected from local residents, hunters and game observers in the area provide the best indirect evidence for determining its status. The survey was usually scheduled from 5 am to 8 am and 4 pm to 9 pm. The survey was taking under the following material; camera, water, battery, rifle, blank page, ballpoint, binocular, hunting Chukar, brown cloths.

**Table 1:** Selected sampling sites in Bajaur District for collection of data on distribution and population of the Chukor Partridges

| Study Zone | Sampling sites | Sub sites |
| --- | --- | --- |
| A | Shingas | 13 |
| B | Serkai | 5 |
| C | Seri Sar | 7 |

The number of calls of Chukor and time spent at each site were recorded. Calling frequency was calculated by using the following formula:

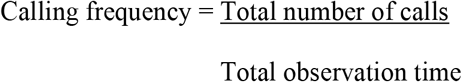

ANOVA (One-way Analysis of Variance) was used to analyze data recorded on call counts during different months of the study period.

## Results

A total of 92 different Chukar signs were recorded at seven sampling points, including 150 direct sightings, 391 calls, 130 feces, 60-foot prints and 190 feathers. Among the seven selected sampling sites, Zulam had the largest species signs (26.09%), followed by Corala (19.57%), and Cicandro (5.43%) had the least signs. Due to the dense vegetation, perhaps due to the sparse population of Chuko (Chukar) in the area, it is difficult to spot birds in the wild. Chukar’s call was also heard in the study area (Table 2). The most calls were recorded at Zulam and Matako sites (80per call), and the fewest calls (two) were recorded at Sikandro site. Difference between the first and second, call was 14-6 seconds during the breeding season, while during non-breeding season, this difference was 50-60 seconds or even more than 2-3 min. In the same way, number of calls per minute (calling frequency) was also the highest as 0.15 during January, 0.3 in February and 0.28 in March indicating the onset of breeding season, while recorded 0.1 in April, 0.07 in May, 0.05 in June, 0.02 in July and 0.01 in August. This was the time when males and females started to pair up. One Way Analysis of Variance (ANOVA) showed a significant difference (df = 8, F 2.90, p < 0.05) in call counts during different months of the year.

**Table 2:** Field signs of *Alectoris chukar* recorded at different sampling sites in District Bajaur during field survey

| Site | Calls | Field sightings | Feathers | Fecal droppings | Foot prints | Total signs | Percentage (%) |
| --- | --- | --- | --- | --- | --- | --- | --- |
| Zulam | 8 | 4 | 7 | 2 | 3 | 24 | 26.09 |
| Matako | 8 | 2 | 2 | 1 | 1 | 14 | 15.22 |
| Kolala | 7 | 4 | 4 | 2 | 1 | 18 | 19.57 |
| Badagai | 5 | 1 | 4 | 3 | 1 | 14 | 15.22 |
| Sikandro | 2 | 1 | 0 | 2 | 0 | 5 | 5.43 |
| Hayaty | 3 | 1 | 2 | 1 | 0 | 7 | 7.61 |
| Shinkot | 6 | 2 | 0 | 2 | 0 | 10 | 10.87 |

Entirety of 181 birds was assessed in 25 areas of the three study zones. In Study Area A, the average total number during the entire study period was estimated to be 60 individuals. However, the winter population (n = 80) was higher than the summer population (n = 39). The population of District A fluctuated in different months of the two seasons. The population was higher in January (n = 87), but declined in February (n = 70), then increased in March and April (n = 81, n = 84), and in summer, from May to August, the population showed a slight growth trend (Table 3). Among the three regions, study area B had the highest population (n = 73), with an average population of 73 in summer and 72 in winter. The highest population in May (n = 80), followed by a decline in June (n = 69) and July (n = 71), but the population in August gradually increased (n = 73). In the different months of winter and summer, the population of this bird in area C was similarly affected. The average number of 54 birds in winter fell to the average number of 48 birds in summer, with the highest in March (n = 56), followed by January (n = 55), February (n = 54) and April (n = 52). Also in the summer, the population in May (n = 34) gradually increased after June (n = 41), July (n = 54), and then declined in August (n = 40) (Table 3). Overall seasonal data shows that there is almost an upward trend, reaching the maximum in January (n = 218). This figure may be due to the reproductive output of this bird. The reason for the decline may be the change of its activities, because the climate has gradually become bad, most zones of the study area are not covered by crops and other vegetation, interns may change the behavior and activities of this bird. During summer and early winter, the bird was also found to be more active and vocal. The low summer populace may be because of the reasons that in this season the bird involved high elevation, which some of the time are distant and can cause inconsistencies in assessing its populace. It is national bird of Pakistan, locally known as zark for male and zarka for female. It can live rocky locations, mountainous in mixed habitat types. Atmosphere is bone-dry to semiarid, temperature shifts and water is commonly accessible from dissipated sources, nest basic scratches lined feather or grass in brushy areas or rocky.

**Table 3:** Population estimation of *Alectoris chukar* in different months during the year 2020

| Zone | Jan | Feb | March | April | Sub. Total | May | June | July | August | Sub. Total | G. Total |
| --- | --- | --- | --- | --- | --- | --- | --- | --- | --- | --- | --- |
|  | Winter |  |  |  |  | Summer |  |  |  |  |  |
| A | 87 | 70 | 81 | 84 | 80 | 35 | 36 | 41 | 43 | 39 | 60 |
| B | 76 | 72 | 69 | 72 | 72 | 80 | 69 | 71 | 73 | 73 | 73 |
| C | 55 | 54 | 56 | 52 | 54 | 34 | 41 | 54 | 40 | 42 | 48 |
| Total | 218 | 196 | 206 | 208 | **207** | 149 | 146 | 166 | 156 | 154 | 181 |

## Discussion

IUCN categorizes Chukor Partridge as the “least concern”, with a stable population trend and a very large population (Birdlife International, 2016). Rich et al. (2004) estimated its global population as c. 2,000,000 people. However, there are reports that its population has decreased from some parts of the world, such as in Europe, where it is estimated that the small population has decreased by nearly 30% in 11.7 years (three generations) (BirdLife International, 2015).

During the investigation, the bird was found to be very active at dawn and dusk, and was found to be engaged in various activities such as sitting, calling, eating and flying during the investigation. We recorded various signs of Chukar in the field, including its calling, feathers, feces, direct sightings and footprints to confirm its occurrence in the study area. The largest overall sign of birds was recorded in Zulam. During the field investigation, it was discovered that the Chukor was related to lower region bushes, including the *Dodona viscosa* as reported by Roberts (1991).

The current survey was conducted in Bajaur district from December 2019 to August 2020. Chukar reproduces once a year according to environmental conditions. Breeding occurs from April to July, with 7 to 21 eggs laid in each season, and an average incubation time of 24 (Christensen, 1996). According to (Del Hoyo et al., 1994; Cole et al., 1995; Christensen, 1996; Perrins, 2003), different results were obtained in this survey (Table 3), since they proposed April to July rearing season for *Alectoris chukar* and the current investigation recorded reproducing season from early March to mid of August. Range eggs per season, 7 to 21 and were gathered 5 to 20 eggs for each season. The most well-known call was a low throw, chuk, chuk utilized by both genders that changes bit by bit to a chukar, chukar and can be gotten with significant distances (Del Hoyo et al., 1994; Cole et al., 1995; Christensen, 1996; Perrins, 2003). Our results showed that the male Chukar in the Bajaur district started to call in January and early February, and then a pairing occurred. The peak breeding season was March and April. Our results on Chukar Partridge’s pairing and reproduction records are consistent with earlier studies by Del Hoyo et al. (1994), Christensen (1996) and Mahmood et al. (2019).

Deforestation, agricultural activities, habitat destruction caused by infrastructure, landslides, and disturbance caused by population growth are the main threats to bird fauna (Khalid et al., 2017). According to reports, the earlier mentioned threat is the main threat to the Chukar partridge population in Bajaur.

## Acknowledgement

We are thankful to all the local peoples who support us throughout our research.

